# SCiMS: Sex Calling in Metagenomic Sequences

**DOI:** 10.64898/2026.02.17.705110

**Authors:** Hanh N. Tran, Kobie J. Kirven, Emily R. Davenport

## Abstract

**Background:** Host sex is a critical determinant of microbial community structure across many host species, influenced by hormonal profiles, physiology, and sex-stratified behaviors. Despite its importance, sex metadata is frequently missing in microbiome studies, including for animal-associated samples. Host chromosomal sex can be inferred from the host-derived reads present in metagenomic data, but existing genomic sex prediction tools rely on fixed coverage thresholds calibrated for human XY chromosomes and require relatively high host reads, limiting their use on low host-biomass samples such as stool and on organisms with other sex-determination systems.

**Results:** Here, we present SCiMS (Sex Calling in Metagenomic Sequences), a bioinformatic tool that leverages host-derived DNA within shotgun metagenomic data to predict host chromosomal sex, even at low host coverage. SCiMS uses a multinomial likelihood computed from observed read counts under each sex and reports chromosomal sex calls. Because the expected read distribution is derived directly from chromosome lengths and ploidy under each candidate karyotype, SCiMS applies to any organism with a heterogametic sex-determination system. We benchmarked SCiMS against existing tools on simulated metagenomic data, human metagenomic samples spanning multiple body sites, and metagenomic samples from seven animal species. SCiMS matched or outperformed existing tools, with its noticeable advantage at low host read conditions.

**Conclusions:** SCiMS provides an accurate, scalable, and cross-species generalizable solution for host chromosomal sex classification, even when host DNA is minimal. By enabling recovery of missing sex metadata, it serves as a quality-control tool analyses in microbiome research. SCiMS is freely available at http://github.com/davenport-lab/SCiMS.

## Introduction

Host chromosomal sex plays a significant role in shaping microbiome composition and function. A multitude of studies have reported sex-specific differences in gut and other microbiota [1–6]. For example, a large human cohort study reported differences between sexes in the relative abundances of major bacterial phyla, including Firmicutes, Actinobacteria, and Bacteroidetes [2, 6]. As a result, recognizing host sex as a key variable in microbiome research is critical to ensure more accurate interpretation of biological patterns in both clinical and ecological contexts.

Despite its importance, host sex information is under-reported in microbiome datasets, particularly for animal associated samples. Our survey of animal associated entries in the AnimalAssociatedMetagenomeDB [7] showed that host sex metadata was missing from 100% of chicken (*Gallus gallus*, n=824), 95.4% of cattle (*Bos taurus*, n=1,336), and 99.9% of pigs (*Sus domesticus*, n=1,121) entries. Even for the common model organism, the house mouse (*Mus musculus*, n=2,280), 88.7% of samples failed to include host sex information. This shortfall could simply be due to researchers not recording sex at the time of sampling, missing entries, only recording sex metadata within supplemental material and not uploading it to public repositories, samples lacking accompanying phenotypic data, or an organism’s sex is not readily determined in the field. Consequently, those aiming to analyze sex-specific patterns or simply control for sex as a confounding factor often find that large portions of otherwise valuable sequence data cannot be used.

Because host-derived reads are routinely present in metagenomic datasets, host chromosomal sex can be in principle inferred directly from sequence data alone even when sex metadata are missing [8]. Several genomic approaches infer sex from whole-genome sequencing data by comparing the proportions of host reads that map to the sex chromosomes, or sex chromosomes relative to the autosomes, including BeXY [9], Rx [10], and Ry [11]. BeXY classifies sex-linked scaffolds by modeling differences in chromosomal ploidy (copy number) across samples using sequencing read counts [9]. Rx calculates the average ratio of reads mapping to X chromosome relative to reads mapping to each autosome [10]. Ry calculates the fraction of reads mapping to Y chromosome relative to reads mapping to both X and Y chromosomes [11]. However, these methods were developed primarily for whole-genome applications in humans, and they share two properties that limit their use on host-derived metagenomic reads. First, most rely on fixed coverage ratio thresholds calibrated for the human X and Y chromosomes, which do not necessarily transfer to organisms with different sex chromosome architecture or to ZW sex-determination systems. Second, Rx and Ry in particular require moderately high host read depth to produce confident calls. This poses a challenge for metagenomic samples where host reads can be sparse, particularly in microbe-rich samples such stool where host-derived reads typically comprise only 1% of total sequences [12]. While the feasibility of calling host chromosomal sex from metagenomic reads has been demonstrated [8, 13], no readily usable tools currently exist that support this task across the range of organisms and sequencing conditions encountered in microbiome studies. Additionally, existing methods have not been systematically benchmarked under realistic metagenomic conditions, including low host read depths and the potential for off-target alignment of microbial reads to the host genome. Together, these limitations motivate the need for methods that reliably infer host chromosomal sex information from metagenomic sequencing data alone, even when host read coverage is minimal.

Here we introduce SCiMS, Sex Calling in Metagenomic Sequences, a command-line tool that determines host chromosomal sex directly from shotgun metagenomic data, even when only a few hundred host reads are present. SCiMS evaluates the multinomial likelihood of the observed chromosome read counts under each candidate karyotype and reports a likelihood-ratio statistic, from which a chromosomal sex is inferred in organisms with heterogametic sex determination systems (XY or ZW). We benchmark SCiMS against BeXY, Rx, and Ry on simulated metagenomic data spanning a range of host read depths and microbial-to-host read ratios, on human metagenomic samples from multiple body sites and populations, and on metagenomic samples from seven animal species. SCiMS provides output as a simple text file of sex calls that can be incorporated into downstream analyses as covariate or used as a metadata quality-control check. Notably, the host-read removal step that is standard in most metagenomic processing pipelines is itself the natural point at which SCiMS can be applied: host reads identified during this step can be passed to SCiMS to recover host chromosomal sex before they are discarded. SCiMS is therefore useful for researchers processing their own human metagenomic data and for animal-associated metagenomic studies.

## Materials and methods

### SCiMS workflow and sex-inference algorithm

SCiMS infers host chromosomal sex by comparing the observed distribution of reads across chromosomes to that would be expected of a male versus a female host. Under each sex hypothesis, the expected fraction of reads on each chromosome is determined by that chromosome’s length and ploidy: a chromosome with two copies contributes twice as much expected read density (i.e. reads mapped per base pair) as one with a single copy. For example, in a female sample (XX), the X chromosome should produce roughly twice the read density of the same chromosome in a male sample (XY), while the Y chromosome should produce no reads under the female hypothesis. SCiMS compares observed counts against these expectations under both hypotheses and reports the more likely sex. Because SCiMS relies on chromosome-wide read density patterns, it should be applied to raw, quality-controlled metagenomic sequencing data prior to host read filtering. Host read removal procedures can distort the relative representation of chromosomes, particularly sex chromosomes, making the remaining reads unsuitable for reliable sex inference.

SCiMS computes the expected read distribution across chromosome under each sex hypothesis, assuming reads are distributed in proportion to each chromosome’s effective size, that is, its length multiplied by its ploidy under that sex. For a host genome containing autosomes and sex chromosomes, the expected probability of observing a read on chromosome c under sex hypothesis S is:

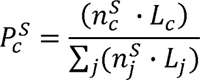

Where *L*_c_ is the length of chromosome c and 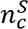 is its ploidy under sex S. Autosome are diploid in both sexes, the homogametic chromosome is diploid in homogametic sex and haploid in the heterogametic sex, and the heterogametic chromosome is haploid in the heterogametic sex and absent in the homogametic sex.

Given observed read counts *N*_c_ across the specified scaffolds and total mapped reads *N* = *∑_c_ N_c_* the log-likelihood ratio comparing male and female hypotheses is computed as follow:

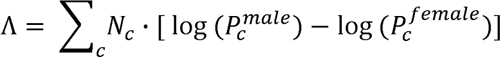

This represents the log-ratio of two multinomial likelihoods. With a uniform prior on sex, the posterior probability that the sample is male is given by the following logistic transformation:

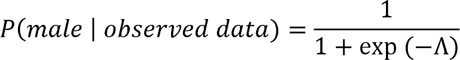

SCiMS reports a confident call when the maximum posterior probability exceeds a user-defined threshold. *τ* (default .*τ* = 0.95): the sample is classified as male if *P*(*male | obserred data*) ≥ *τ*, female if *P(female | obserred data)* ≥ *τ*, and uncertain otherwise.

The complete mathematical framework, including multinomial likelihood derivation and a discussion of mipmapping rate is provided in Supplementary Methods.

### Comparative method performance

To benchmark SCiMS, we directly compared its performance against three existing sex inference tools originally developed for genomic data: BeXY, Rx, and Ry. We applied all methods to the same mapping statistics (.idxstats) input files to ensure a consistent basis for comparison. We evaluated each method using standard classification metrics: accuracy, precision, recall, and F1 score, calculated separately for each sex as following:

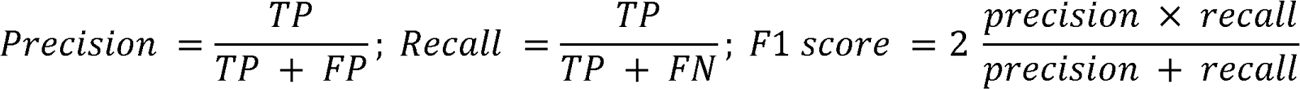

where TP (true positives) denotes correct predictions, FP (false positive) refers to incorrect predictions, and FN (false negatives) represents cases where the method is abstained (i.e., uncertain calls).

### Simulated dataset

Metagenomic samples often contain a low proportion of host-derived reads. To evaluate how well SCiMS perform under such conditions, we simulated sequencing data across a wide range of host read depths. Specifically, we generated samples by combining host-derived reads with microbial reads from a published microbial community simulation: CAMI II [14]. This design captures two features of real metagenomes: (i) microbial reads that can spuriously align to the host reference genome, and (ii) variable host fractions across the range observed in real metagenomic studies.

*1. Host read generation.* We constructed a male and a female human reference genome from the genome assembly GRCh38.p14 using a custom script. The male reference included a full diploid genome (autosomes plus both X and Y chromosomes), while the female reference retained all autosomes along with two copies of the X chromosome. From each reference genome, we simulated 100 million paired-end reads (150 bp, 1% sequencing error rate) using wgsim v.0.3.1-r13 [14]. These reads constitute the host-only read pools from which sample-level read subsamples are drawn. To create individual replicates, we subsampled the host read pools using seqtk v.1.4 [15] with a unique random seed per replicate. For each combination of sex (male, female), host read depth (150, 250, 350, 450, 1,000, 10,000 reads), and replicate (1– 100), we generated paired-end FASTQ files containing the exact host read pairs. This produced 1,400 unique host-only samples covering a wide range of host read depths.
*2. Source of microbial reads.* Microbial reads were drawn from the Critical Assessment of Metagenome Interpretation (CAMI) II, sample S004 of the 450-genome oral metagenome simulation [14]. This dataset provides a realistic representation of a complex microbial community of known composition. We treated this as our microbial read pool and subsampled from it without replacement using seqtk v.1.4 [15].
*3. Metagenomic sample generation.* For each host sample described above, we generated mixed metagenomic samples by concatenating the host reads with a defined number of microbial reads from the CAMI pool, controlled by a host fraction parameter *f* ∈ {0.001, 0.01, 0.1, 0.5, 0.95, 1.0}. Given a host depth *h* ∈ {150, 250, 350, 450, 1,000, 10,000} and host fraction f, the number of microbial reads added is 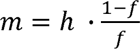, so that the host reads constitute exactly fraction f of the final mixed sample. For example, a sample with 10,000 host reads at a host fraction of 0.001 receives 9,990,000 microbial reads, yielding a metagenomic sample of 10,000,000 reads in which host reads make up 0.1% of the total reads. The mixing is performed at the FASTQ level prior to alignment, reproducing the alignment conditions encountered in real metagenomic sequencing datasets. This captures the potential for off-target alignment of microbial reads to the host reference genome under realistic metagenomic processing conditions. The resulting design comprises 2 sexes × 6 host depths × 6 host fractions × 100 replicates = 7,200 simulated metagenomic samples.
*4. Downstream alignment and sex inference.* Mixed FASTQ files were processed through the same alignment pipeline used for real metagenomic datasets (Materials and Methods, “Data preprocessing”): mapping to GRCh38.p14 with Bowtie2 v.2.5.2 [16], retain uniquely mapped reads with MAPQ ≥ 30 using SAMtools v.1.19 [17]. Chromosome-level alignment statistics were extracted with SAMtools idxstats and used as input for SCiMS, BeXY [9], Rx [10], and Ry [11]. Sex calls were compared against the known sex labels for each simulated sample.
*5. T2T-CHM13v2.0 simulation.* To assess whether the choice of host reference assembly affects classification outcome, we repeated the simulation benchmark using the T2T-CHM13v2.0 assembly (NCBI RefSeq GCF_009914755.1). Host reads were generated similar to those in the GRCh38 simulation (section 1) and mixed with microbial reads from the CAMI II metagenomic sample across the same host fractions and host read depths. Reads were aligned to the T2T-CHM13v2.0 reference and processed similarly to the GRCh38 simulation. Classification outcomes from the two assemblies were compared at matched host read depth and host fraction.

### Real metagenomic datasets

To determine whether simulated performance translated to real metagenomic data, we applied SCiMS to ten real metagenomic datasets **(Table 1).**

**Table 1.** Metagenomic datasets. (*) species lacking a chromosome-level reference assembly with annotated sex chromosomes, and reads were mapped to the reference genome of a closely related species.

| Dataset | Species | Samples (n) | Sex | Accession / Citation |
| --- | --- | --- | --- | --- |
| Human Microbiome Project | *Homo sapiens* | 1339 | 718 males, 621 females | dbGaP phs000228 [12] |
| Hadza Hunter–Gatherers | *Homo sapiens* | 331 | 169 males, 162 females | PRJEB49206 [18] |
| Indian Human Gut Microbiome | *Homo sapiens* | 99 | 49 males, 50 females | PRJNA397112 [19] |
| Baboon | *Papio cynocephalus* * | 48 | 17 males, 31 females | PRJNA271618 [20] |
| Cow | *Bos taurus* | 47 | 47 males | PRJEB31266 [21] |
| Pig | *Sus scrofa* | 112 | 40 males, 71 females | PRJNA629856 [22] |
| Black rhino | *Diceros bicornis* * | 20 | 11 males, 9 females | PRJNA532626 [23] |
| Mesquite lizard | *Sceloporus grammicus* * | 20 | 20 males | PRJNA1106546 [24] |
| Mouse gut | *Mus musculus* | 111 | 23 males, 88 females | ENA PRJEB32890 [16] |
| Chicken cecum | *Gallus gallus* | 94 | 22 males, 72 females | ENA PRJEB33338 [25], PRJNA658643 [26], & PRJNA947933 [27] |

### Data preprocessing

For simulated samples, we directly aligned the reads to human GRCh38.p14 assembly using Bowtie2 v.2.5.2 [16]. For real metagenomic datasets, we trimmed adapters and filtered reads for quality using fastp v.0.23.4 [28] with default parameters. For all datasets, we used only samples with at least 100 host reads. We then mapped the filtered reads to the corresponding host reference genomes (human GRCh38.p14, mouse GRCm39, and chicken GCA_027557775.1) using Bowtie2 v.2.5.2 [16]. After alignment, we removed potential PCR duplicates with the MarkDuplicates function in Picard Tools v.3.1.1 (http://broadinstitute.github.io/picard/). We further refined the alignments by retaining only uniquely mapped reads with mapping quality (MAPQ) ≥ 30, using SAMtools v.1.19 [17]. To avoid confounding effects from pseudoautosomal regions (PARs), which are present on both X and Y chromosomes and share high sequence similarity [29], we explicitly excluded these regions from the human dataset using BEDtools v.2.31.0 [30]. Finally, we used SAMtools’ function idxstats to extract the number of reads mapped to each chromosome or scaffold, generating per-sample read count profiles that served as the primary input for SCiMS.

## Results

### Overview of SCiMS

We developed SCiMS, a command-line tool designed to recover host chromosomal sex directly from metagenomic alignments. Unlike methods relying on fixed thresholds for sexing, SCiMS uses per-chromosome read counts as a multinomial distribution and compares the observed distribution against the ploidy-weighted expectation for each sex. Specifically, for a given chromosome *c* under sex *s*, the expected probability of a read coming from *c* is proportional to its ploidy 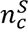 (2 for autosomes; 2 or 1 for the homogametic chromosomes; 1 or 0 for the heterogametic chromosome) times its length *L_c_*, normalized over all chromosomes considered. A likelihood-ratio test across these per-chromosome counts yields the log-odds of male versus female, which is converted to a posterior probability under a uniform prior. To remain robust to the spurious cross-specifies alignments characteristic of metagenomic data, where a single misaligned microbial read against a zero-ploidy chromosome would otherwise force an infinite likelihood ration, SCiMS applies a small mismapping constant *ε* = 10_—4_ to expected probabilities before evaluating the test. Because the model is analytical and parameter free, it generalizes across any heterogametic species (XY or ZW) and any reference genome. Full mathematical detail is provided in Supplementary Methods.

The recommended workflow begins with preprocessing of raw sequencing reads (**Fig. 1**). First, users align reads to a host reference genome using an aligner such as Bowtie2 [16]. To improve classification accuracy, we recommend removing PCR duplicates with Picard, then sorting and filtering the BAM files using SAMtools to retain only high-quality, uniquely mapped reads (MAPQ ≥ 30). When annotations are available, filtering out pseudoautosomal regions (PARS) helps avoid coverage biases from regions shared between sex chromosomes [29]. SCiMS accepts either a mapping statistic file (.idxstats) generated by SAMtools or a BAM file which is convenient for users who already align reads to a host reference as part of host read removal prior to microbial profiling.

**Fig. 1.**
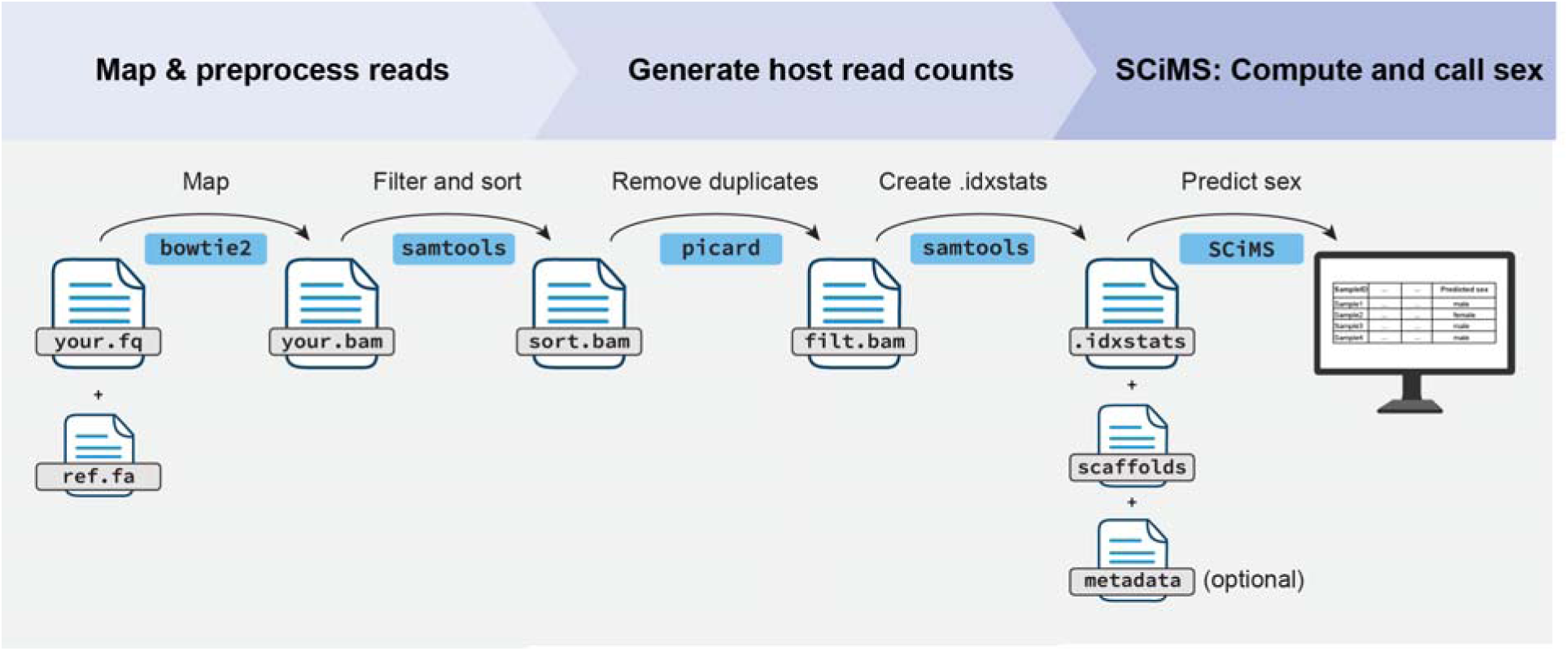
Suggested workflow. Shotgun metagenomic reads (your.fq) should first be aligned to the host reference genome (ref.fa) with a mapping tool (such as Bowtie2) to produce a BAM file (your.bam). The alignment should then be sorted and filtered for mapping quality with SAMtools (sort.bam), after which PCR duplicates should be removed with Picard or similar (filt.bam). SCiMS takes input as either the mapping statistics file generated by SAMtools (.idxstats) which supplies read counts per chromosome or the sorted BAM file, together with a text list of chromosome or scaffold identifiers from the host reference genome, and optionally, a metadata table. SCiMS evaluates a multinomial likelihood-ratio test comparing observed read counts against expected distribution for each sex and returns a probabilistic chromosomal sex assignment per sample displayed in a tabular summary and amended to the metadata table if supplied.

SCiMS supports both XY and ZW sex-determination systems through the ‘--ZW’ option, extending its applicability to a broad range of organisms with heterogametic sex chromosomes. The classification threshold on the posterior probability defaults to 0.95; users seeking higher sensitivity can lower this threshold, while those needing stricter calls can raise it toward 1. A complete tutorial and implementation guide is available at (https://github.com/davenport-lab/SCiMS).

### SCiMS classifies host chromosomal sex accurately at low host read depths

To quantify classification accuracy directly, we benchmarked SCiMS against BeXY, Rx, and Ry on simulated metagenomes with known host sex. Each sample was generated by combining simulated host reads with microbial reads from the CAMI II oral metagenomic simulation [14] at defined ratios. Because the true karyotype is fixed by construction, these simulations isolate the effect of host read depth and microbial dosage on inference accuracy, variables that cannot be controlled in empirical data. We generated 7,200 samples spanning a broad range of host read counts (150 to 10,000 reads) and six host-to-microbial read fractions, from host-only (100% host) to predominantly microbial (0.1% host; Materials and Methods).

As expected, all four methods classified host sex almost perfectly when host reads were abundant (>10,000 reads; **Fig. 2a**), and the methods were therefore distinguished primarily by their behavior at low host read depth. Specifically, SCiMS and BeXY were broadly comparable and both substantially exceeded Rx and Ry. Overall accuracy for SCiMS and BeXY rose from 0.52 and 0.53 at 150 host reads to 0.97 and 0.99 by 1,000 reads (**Fig. 2a**), whereas Rx and Ry remained considerably lower across the same range. The difference between these two groups of methods reflects their use of the available signal. SCiMS and BeXY evaluate the joint distribution of reads across chromosomes, whereas Rx and Ry rely on a single ratio and abstain when coverage of the relevant chromosome is insufficient. SCiMS and BeXY, nonetheless, differed in a sex-specific manner (**Fig. 2b–c**). SCiMS achieved higher accuracy on female than on male samples at low read depths, while BeXY achieved slightly higher accuracy on male samples. These patterns were reflected in per-class recall (**Fig. 2c**). SCiMS recovered more female (0.89) than male (0.77), consistent with a tendency of abstain when the heterogametic chromosome is sparsely covered, whereas BeXY showed the opposite, with marginally higher male (0.82) than female (0.80) recall. Rx recovered relatively few samples of either sex (male recall 0.55, female recall 0.62). Ry exhibited the most noticeable asymmetry, with perfect female recall (1.00) but low male recall (0.26). This behavior follows directly from its decision rule, which treats the absence of Y-mapped reads as evidence of female sex, which recover females reliably but misclassifies males as females when host reads, and thus Y-mapped reads, are scarce.

**Fig. 2.**
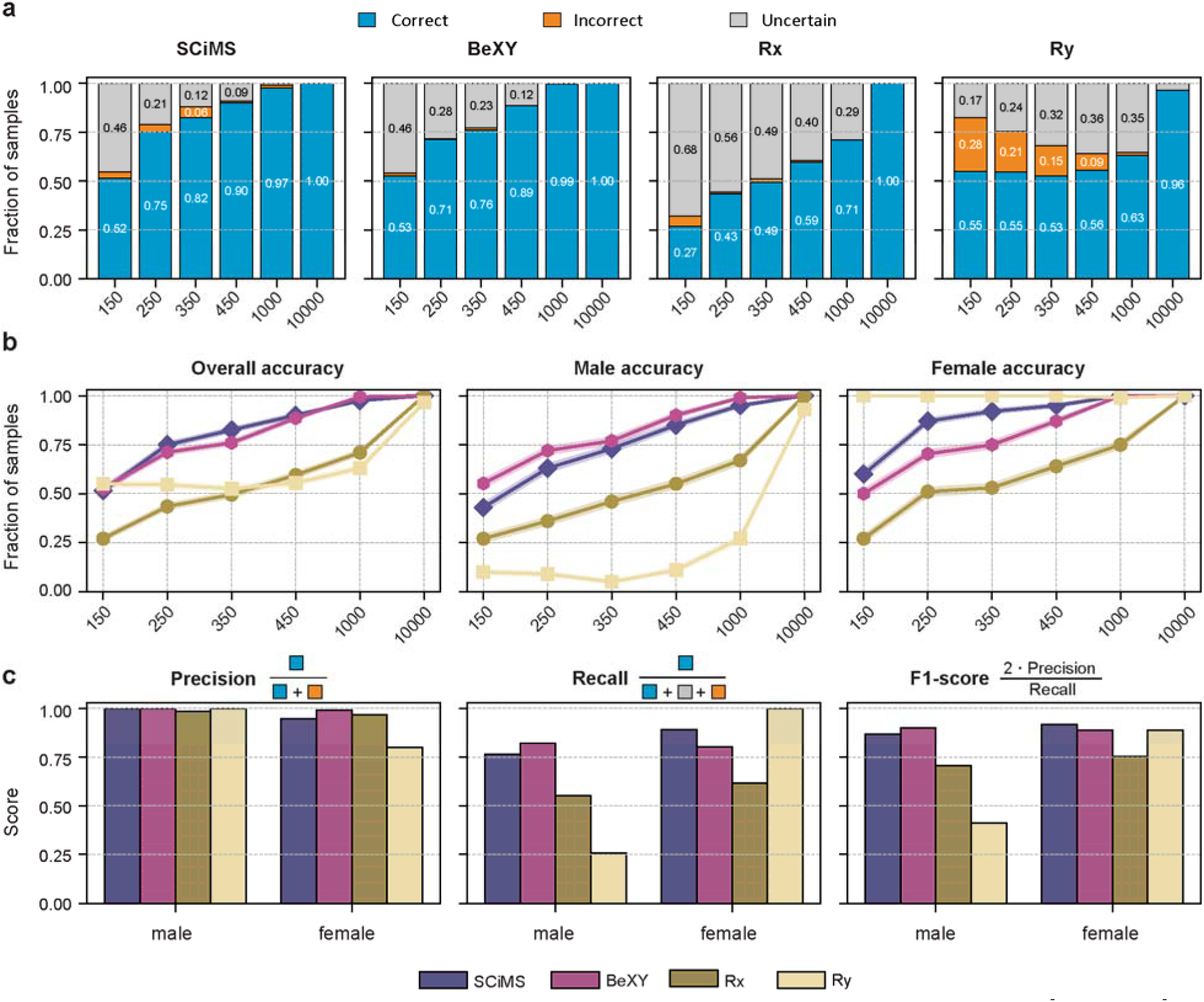
Benchmarking SCiMS, BeXY, Rx, and Ry on simulated metagenomic samples (n=8,400). Host-derived reads were combined with microbial reads from the CAMI II oral metagenomic simulation across host-to-microbial concentrations and sex host reads depth (150–10,000 reads), yielding 7,200 simulated samples with known sex (Materials and Methods) **a)** Fraction of samples classified as correct (blue), incorrect (orange), and uncertain (grey) by each method at each host read depth. **b).** Call accuracy, defined as the fraction of confident calls that were correct, as a function of host read depth, shown for all samples (left panel), male samples (middle panel), and female samples (right panel). Lines show mean accuracy across replicates and shaded bands indicate ±1 standard error of the mean. **c)** Precision, recall, and F1 scores for each method, computed per sample and averaged across all samples at all read depths. Precision is defined as correct / (correct + incorrect), recall as correct / (correct + incorrect + uncertain), and F1 as their harmonic mean.

To test whether microbial reads mis-mapping to the host genome affects classification, we examined accuracy as a function of proportion of host reads in each sample while holding host read depth fixed. For all methods, accuracy depended on the number of host reads available but was unchanged across host fractions at the same depth (**Supplementary Fig. 1S**), indicating that the microbial background did not measurably bias sex calls even when host reads constituted as little as 0.1% of the sample. Additionally, to assess whether the choice and completeness of the host reference genome affect classification, we repeated the simulation benchmark using two human assemblies: the standard GRCh38.p14 reference and the T2T-CHM13 assembly (**Supplementary Fig. S1a-b**). For all four methods, accuracy was determined primarily by host read depth and was largely insensitive to the reference used. Specifically, the relative ordering of the methods was unchanged, and most methods reached comparable accuracy on both assemblies. The exception was Ry, which showed poor recall across read depths in the T2T-CHM13 simulations, particularly for male samples. We hypothesize that this reflects the increased repetitive content represented in the completed T2T Y chromosome assembly, which reduces the number of uniquely mappable Y-derived reads available for sex inference. Because Ry relies directly on detection of Y-chromosome reads, it may be particularly sensitive to this reduction in informative reads. Within each reference, accuracy was likewise stable across host fractions at the same read depth, from host only samples to samples in which host reads constituted as little as 0.1% of the total reads. Together, these results indicate that host read depth is the primary determinant of classification accuracy across the range of host-to-microbial read ratios and reference genomes examined here.

Taken together, results from simulated data indicate that SCiMS and BeXY are the most robust to low host read depth, with complementary sex-specific behavior, while the ratio-based methods Rx and Ry are limited by their reliance on a single chromosome and their resulting tendency to misclassify when that chromosome is poorly covered.

### SCiMS achieves accurate and reliable host sex classification across human metagenomic datasets

To evaluate performance on real data, we applied all four methods to 1,727 human metagenomic samples with host sex recorded in the associated metadata. The dataset includes three human cohorts: the Human Microbiome Project [12], together with fecal samples from non-Western populations including the Hadza hunter-gatherers [18] and Indians [19] **(Supplementary Table 1).** These cohorts span a wide range of host DNA abundance, from host-rich samples (vaginal and anterior nares) to highly microbial-skewed samples (stool). To capture how each tool performs across the depth range, we stratified samples by host-aligned read count (<500, 500–1,000, 1,000–10,000, and >10,000) and evaluated both accuracy (correct calls / total called samples) and call rate (called samples / total samples) (**Fig. 3**).

**Fig. 3.**
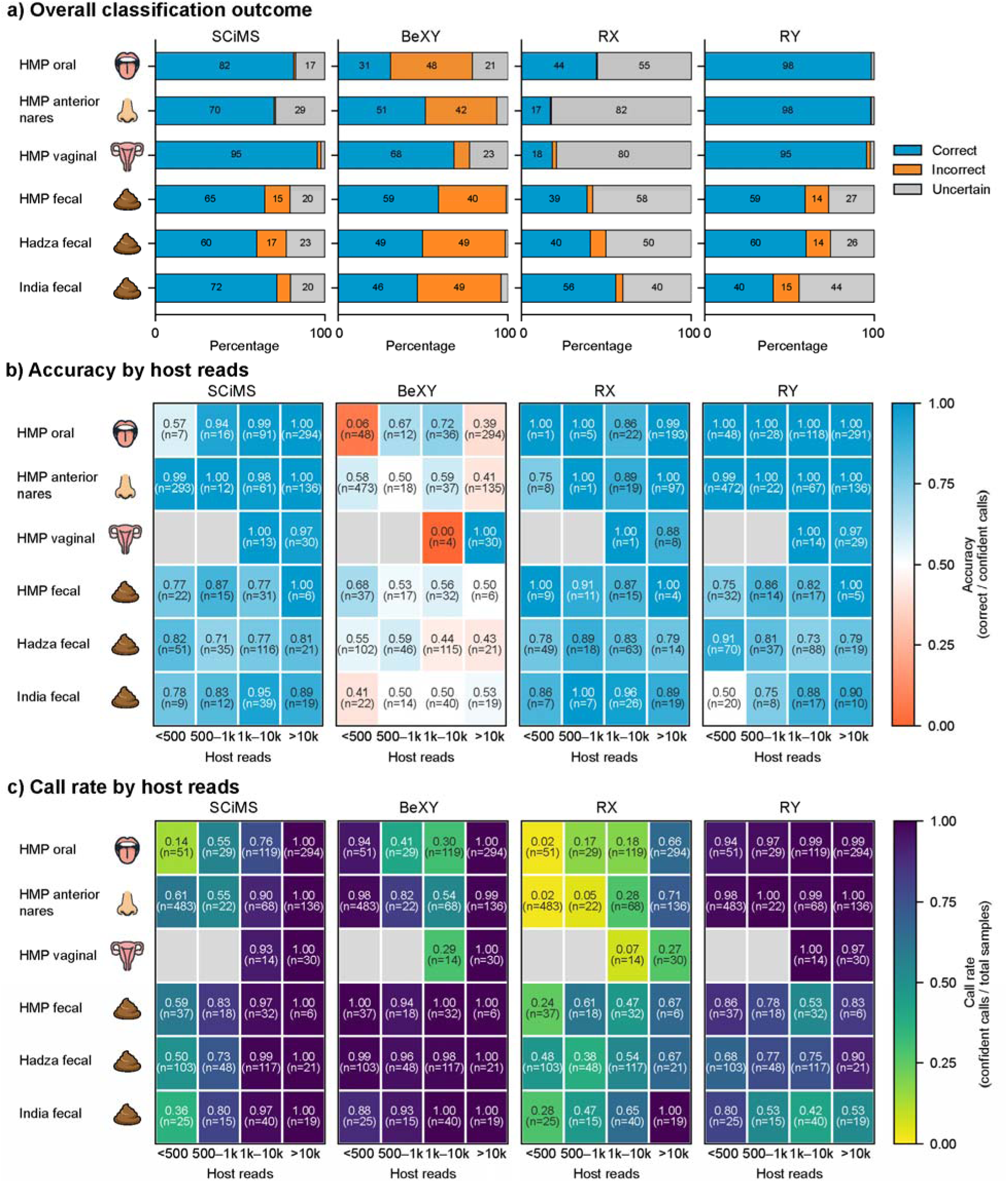
Sex-classification performance across real human metagenomic cohorts spanning a wide range of host sampling sites. SCiMS, BeXY, Rx, and Ry were evaluated on six human datasets: oral, anterior nares, vaginal, and three fecal datasets. **a)** Breakdown percentage of samples called correctly (blue), incorrectly (orange), and uncertain (gray) by each tool in each dataset, pooled across host read depth. **b)** Classification accuracy (fraction of confident calls that were correct). **c)** Confident call rate (fraction of samples receiving a confident call). In (b) and (c), the value and sample size (n) are shown in each cell. Gray cells indicate no samples in that cohort and read depth combination.

Overall, SCiMS produced correct calls for the majority of samples in every cohort, calling 60–95% of samples correctly while keeping incorrect calls low (.17%; **Fig. 3a**). Similar to simulation results, SCiMS accuracy in real metagenomic samples improved with host read depth across all cohorts (**Fig. 3b**). Specifically, in HMP anterior nares, HMP oral, and HMP vaginal samples, SCiMS accuracy reached 0.88, 0.94, and 1.00 respectively at 1,000 host reads and above. In the more challenging fecal samples (HMP, Hadza, and Indian), where host DNA typically constitutes less than 1% of sequencing reads [12], SCiMS accuracy rose with host read depth in all three cohorts, from 0.71–0.87 below 10,000 host reads to 0.81–1.00 above 10,000 reads (**Fig. 3b**). In simulations, male samples had lower call accuracies at the lowest read depths. This is expected, as at a fixed host read depth, the shorter Y chromosome would be expected to contribute fewer reads than the longer X chromosome. This pattern was not consistently observed in the real metagenomic cohorts (**Supplementary Fig. 2**), where additional sources of biological and technological variation likely outweigh the modest theoretical advantage observed for female samples in simulations.

SCiMS confident-call rate likewise increased with host read depth (**Fig. 3c**). Once host depth exceeded 1,000 reads, SCiMS confidently made calls on 97–100% of samples in every cohort. At lower depths (<500 and 500–1000 reads), where the call rate was lowest, the uncalled samples were disproportionately female. Female call rate was as low as 0.30 in anterior nares and 0.61–0.63 in the oral and Hadza fecal samples, versus ≥ 0.94 for males (**Supplementary Fig. 2**). Notably, this asymmetry was driven by abstention rather than misclassification. Of the 817 female samples in the pooled dataset, SCiMS withheld a confident call on 368 samples (45%) but committed to an incorrectly call on only 44 (5%) samples. Conversely, in the fecal samples, a comparably high call rate was accompanied by lower accuracy. At 1,000–10,000 host reads, for example, SCiMS committed to 97–99% of fecal samples yet classified only 77–95% of them correctly (**Fig. 3b–c**), and overall accuracy in fecal samples rose only modestly with depth, from 71–87% in samples with below 10,000 reads to 81–100% in samples with 10,000 reads and above. This pattern reflects a fundamental limitation of metagenomic sex inference in highly microbial-rich samples (i.e. stools): even when sufficient host reads are present to support a confident call, they may carry too few sex-chromosome reads to resolve sex reliably.

The benchmark tools showed different behaviors in the trade-off between call rate and accuracy. BeXY committed to confident calls on most samples (call rate >0.90 in most groups) but those calls were frequently incorrect. Specifically, BeXY committed to confident calls on most samples of both sexes (call rate ≥ 0.82 for males and ≥ 0.76 for females), but those calls were frequently incorrect, with accuracy falling as low as 0.06 (HMP oral, <500 reads) and remained largely in the 0.40–0.60 accuracy range (**Fig 3a–b**). This manifested as confident misclassification of one entire sex. For example, male accuracy fell to 0.00 in the HMP and Hadza fecal cohorts, and female accuracy to 0.21 in anterior nares and 0.00 in the Indian fecal cohort (**Supplementary Fig. 2**). Rx showed the inverse pattern where the confident accuracy was ≥0.86 in most groups (**Fig. 3b**), but committed to very few samples at low depth, with call rates as low as 0.2–0.28 in oral and anterior nares samples below 10,000 host reads (**Fig. 3c**). This behavior reflects Rx’s algorithm design where it returns uncertain when the confidence interval is too wide to determine sex, committing to only a small fraction of samples. As a result, although its calls were often accurate, its call rate in all sample groups was correspondingly low (as low as 17.1% in anterior nares). Ry achieved high accuracy and call rate in the HMP oral anterior nares, and vaginal samples (≥0.97 across all depths), but accuracy and call rate dropped in the three fecal datasets (**Fig. 3a–b**). This drop was specific to males whose accuracy fell to 0.33 in HMP fecal and 0.12 in the Indian cohort, while female accuracy remained high throughout ((≥0.86, **Supplementary Fig. 2**).

Together, the human results reinforce and extend our simulation findings, establishing SCiMS as a reliable tool for inferring host chromosomal sex across heterogeneous conditions. In simulated data, SCiMS and BeXY were the two most robust methods at low host read depth. In real human metagenomic data, however, BeXY’s accuracy dropped significantly while Ry emerged as the strongest tool, performing well wherever host DNA, and thus Y-chromosome signal in males, was abundant. The leading alternative differed between settings underscores that each existing tool works well only under specific conditions, whereas SCiMS remains robust across both simulated and real data across the full range of host read depths, body sites, and populations examined. It classifies samples accurately and returns ‘uncertain’ more frequently than erring when it could not. This consistency distinguishes SCiMS for general use across conditions encountered in microbiome studies.

### SCiMS accurately classifies host chromosomal sex across diverse animal species, including a ZW sex-determination system

We next applied all four methods to metagenomic samples from seven animal species spanning a range of sex-chromosome systems and reference-genome quality: mouse, cow, baboon, black rhino, pig, mesquite lizard, and chicken (**Table 1**, **Fig. 4, Supplementary Table 2**). In the three species with well-characterized, conspecific XY references, mouse, pig, and cow, SCiMS classified nearly all samples correctly (1.00, 0.98, and 0.85 respectively), as did BeXY (0.96, 0.99, and 0.83 respectively), and Rx (0.86, 0.97, and 0.97 respectively) (**Fig. 4**). Ry classified female samples accurately but correctly called few or no male samples, resulting to its low overall accuracy (0.79 mouse, 0.61 pig, and 0.02 cow) (**Fig. 4; Supplementary Fig. 3**). We further examined three animal species (baboon, black rhino, and mesquite lizard) that lacked a sex-chromosome-resolved conspecific assembly and were instead mapped to a closely related species whose have a complete genome assembly. Reference completeness had limited effect across most of this group. Baboon samples (*Papio cynocephalus*) mapped to the olive baboon (*Papio anubis*) reference were classified as accurately as the conspecific-reference species (0.98 for SCiMS, 0.97 for BeXY, and 0.94 for Rx; **Fig. 4**). Mesquite lizard (*Sceloporus grammicus*) samples were mapped to a green anole lizard (*Anolis carolinensis*) and were correctly inferred up to 0.85 by SCiMS, 0.80 by BeXY, and 0.75 by Rx. Black rhino (*Diceros bicornis*) samples were mapped to Southern white rhino (*Ceratotherium simum simum*) genome assembly. All methods performed relatively worse compared to other cross-mapping species such as baboon and mesquite lizard. Specifically, SCiMS accurately classified 65% of samples, 55% for BeXY, 60% for Rx, and 35% for Ry. Notably, SCiMS was the only method to classify both sexes with comparable performance, male F1=0.70 and female F1=0.63 (**Fig. 4**). BeXY and RY each failed entirely on one sex, BeXY recovering no females and Ry no males, while Rx, though functional on both, was less balanced (male F1=0.74; female F1=0.56). Together, these results indicate that SCiMS remains robust in organisms with incomplete or related-species genome reference. This is consistent with our simulation results in human samples, where SCiMS performed comparably on the GRCh38 and T2T-CHM13 assemblies, indicating that its accuracy is robust to the choice of reference more broadly, whether across reference builds of the same species or across closely related species.

**Fig. 4.**
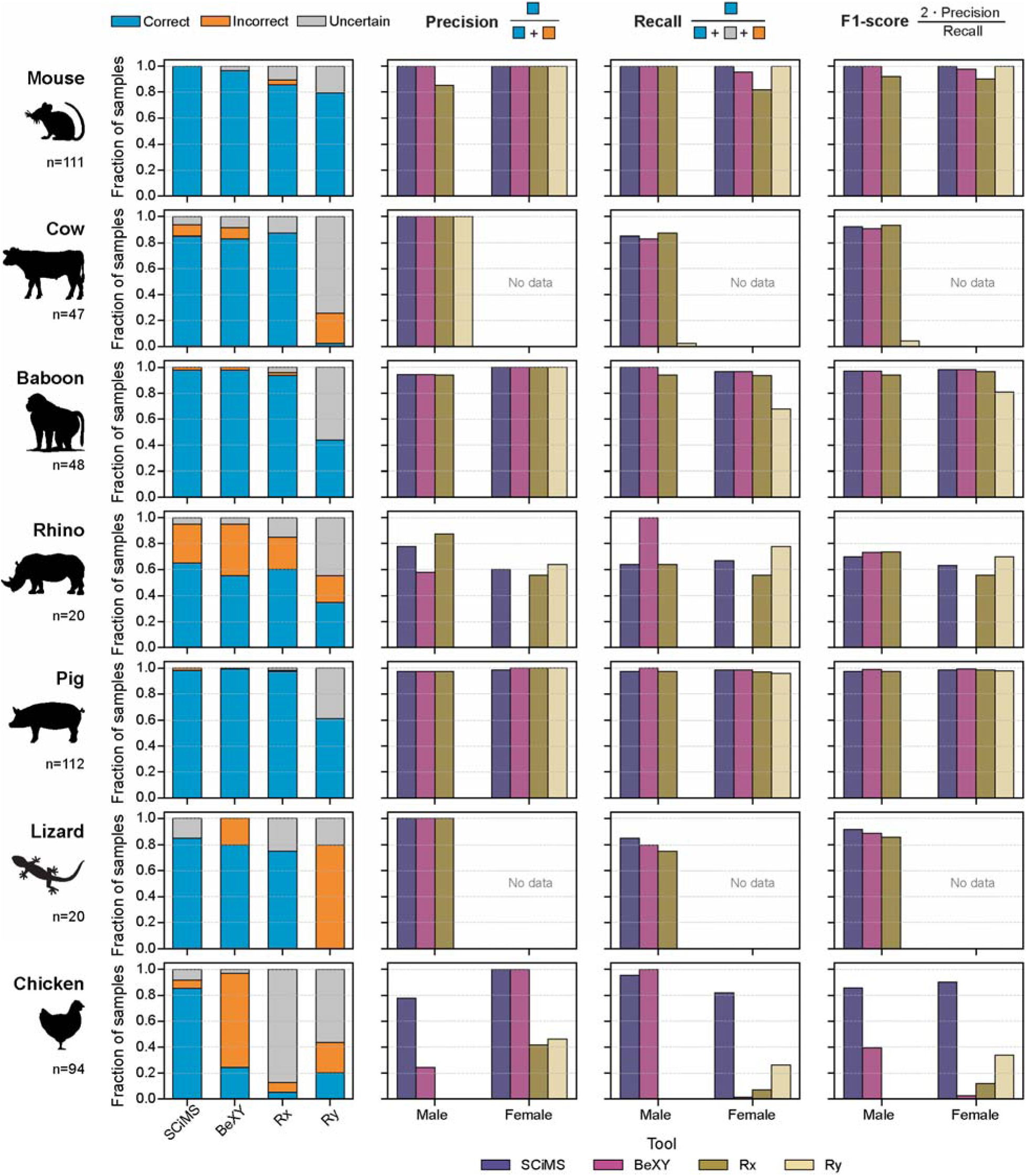
Sex classification performance of SCiMS, BeXY, Rx, and Ry on metagenomic samples from seven animal species: mouse, cow, baboon, black rhino, pig, mesquite lizard, and chicken. Each row corresponds to one animal species, spanning XY and ZW sex-determination systems and conspecific and related species assemblies (Table 1). **(Left column)** fraction of samples classified as correct (blue), incorrect (orange), or uncertain (grey) by each method. **(Right column)** Precision, recall, and F1 score for each method, computed separately for male and female hosts. Precision is the fraction of confident calls of a given sex that were correct, recall is the fraction of true samples of that sex that were correctly called, and F1 is the harmonic mean of precision and recall. Metrics were computed per species and shown by host sex. “No data” denotes species for which a metric could not be computed because the cohort contained only one sex. Absent bars indicate a method made no confident calls of that sex.

To further examine the tool’s broad applicability, we benchmarked SCiMS, BeXY, Rx, and Ry to a chicken cohort, a species with a ZW sex determination system. SCiMS classified 85% of samples correctly, whereas the other three methods largely failed, BeXY classified 24% of samples correctly, Rx 5%, and Ry 20% (**Fig. 4**). Performance metrics stratified by sex showed that SCiMS was the only method to classify both sexes correctly (male F1=0.86; female F1=0.90), while BeXY misclassified nearly all females (male F1=0.39; female F1=0.03), Rx and Ry misclassified nearly all males (male F1=0.00; female F1=0.12, and male F1=0.00; female F1=0.34, respectively).

Together, these cross-species results highlight two methodological advantages of SCiMS design. First, because SCiMS operates on per-chromosome read counts modeled against the expected distribution under each candidate karyotype, it adapts to any heterogametic sex-determination system without retraining or optimization of species-specific parameters. Users specify which chromosomes correspond to the homogametic and heterogametic sex for the target organisms, and the multinomial likelihood ratio is computed accordingly. Tools designed around XY systems (Rx, Ry, and BeXY) failed to accommodate the reversed ZW configuration in chicken. Second, SCiMS was robust to the completeness and choice of reference assembly. It classified host sex accurately when reads were mapped to a congeneric reference. This robustness is consistent with our human simulations, in which SCiMS performed comparably on the GRCh38 and T2T-CHM13 assemblies. Overall, the flexibility across sex-determination systems and tolerance of incomplete or related-species references make SCiMS applicable for inferring host chromosomal sex in diverse non-model organisms.

## Discussion

SCiMS is a command-line tool for inferring host chromosomal sex from metagenomic sequencing data, designed to operate on a per-sample basis without training data or species-specific calibration. The ability to function effectively at low host sequencing depths fills a critical gap in metagenomic analysis, because host reads can be sparse in metagenomic data depending on the site of sampling and unevenly distributed across the genome. By evaluating the multinomial likelihood of the observed host read counts under each candidate karyotype, SCiMS generalizes across heterogametic sex-determination systems (XY or ZW). Because the expected per-chromosome read distribution is computed directly from the input chromosome lengths and ploidy assignments, no species-specific thresholds or recalibration are required. This distinguishes from summary-statistic methods (Rx, Ry) whose decision thresholds are calibrated from human X and Y chromosomes, and cohort-conditional methods (BeXY), which require a batch of samples to fit their posterior model.

Across simulated and real data, SCiMS maintained a balance between making calls and avoiding errors. When host coverage was insufficient to resolve sex, SCiMS was more likely to return ‘uncertain’ rather than committing to incorrect classification. While SCiMS showed similar performance to BeXY under simulated conditions, it consistently outperformed BeXY across the empirical human and animal metagenomic datasets. One possible explanation for this difference is that BeXY was developed to infer sex from chromosome-wide coverage patterns in low-coverage genomic data, whereas real metagenomic samples contain additional sources of variation that may alter chromosome-wide read distributions, including uneven host DNA recovery, microbial background reads, and cohort-specific mapping characteristics. The implications of SCiMS’s performance are significant, and it has a wide range of use cases across fields. First, in biomedical research metagenomic data is increasingly being collected for studying the microbiome’s role in health and disease. Accurate sex determination can be crucial for understanding sex-specific disease manifestations and tailoring treatments accordingly. We have benchmarked SCiMS on two of the most commonly used organisms in biomedical research to demonstrate its performance: Humans and mice. Second, SCiMS will be useful for studies of livestock and wildlife microbiomes, where host sex information may not be available. This could occur in cases where fecal samples are collected from the field without a behavioral observation accompanying collection or from species that demonstrate low sexual dimorphism, making it challenging to sex an organism from a distance or without tranquilizing the animal. The repository for SCiMS includes instructions on how to run SCiMS on new, user-defined reference assemblies so long as the organism has a heterogametic sex determination system.

Finally, SCiMS can also serve as a valuable quality control (QC) step in metagenomic analyses even when sex information has been collected, identifying potential sample swaps or metadata inconsistencies.

However, SCiMS has some limitations and areas that could be further developed or benchmarked. Its accuracy depends on the presence of reads mapping to host sex chromosomes. While it shows high performance at low host read depths, it will still report inconclusive sex calls for samples with extremely low read depths or where host sex chromosome reads are absent. We benchmarked SCiMS performance using a robust posterior probability threshold of 0.95, which was selected to balance precision and recall. However, users wishing for even higher precision have the option to increase that threshold. Additionally, since sex resolution requires sufficient sex-chromosome signal, in the bacteria-rich samples, such as stool, a confident chromosomal sex call could still be unreliable when too few reads mapped to the sex chromosomes. Accuracy in these samples was lower for all methods. Next, while we benchmarked SCiMS on a diverse set of host species, including several cases where only a closely related reference genome was available, performance varied among taxa. SCiMS relies on accurate representation of chromosome structure within the reference genome, and its performance may decline when the reference is highly divergent from the organism being analyzed. For optimal performance, users should benchmark SCiMS on their focal organism and reference genome whenever possible. This consideration may be particularly relevant for applications involving ancient DNA or highly divergent populations, where structural variation and long-term evolutionary change may result in chromosome architectures that differ substantially from those represented in modern reference assemblies. Finally, SCiMS is currently tailored specifically for XY and ZW sex determination systems. This currently prohibits its use in organisms with other sex determination mechanisms, such as those with genetic balance (e.g. *Drosophila*), polygenetic (e.g. zebrafish), or environmental sex determination systems (e.g. alligators and crocodiles, most turtles, and some worms). Regardless, SCiMS offers a novel, accurate new framework for calling sex from host-derived metagenomic samples.

Importantly, the purpose of SCiMS is to infer specifically the chromosomal sex of host organisms with heterogametic sex chromosomes (XY or ZW). Gender identity cannot be inferred using SCiMS, as gender is a social construct not inherently tied to biological sex [31]. Additionally, SCiMS may have limitations in accurately identifying sex in individuals with sex chromosome aneuploidies, where individuals are born with more or fewer than two sex chromosomes. These variations include conditions like Klinefelter syndrome (XXY), Turner syndrome (X), or XYY syndrome (XYY) [32]. Similarly, SCiMS may have limitations in accurately identifying sex in individuals with sex chromosome mosaicism, where two or more genetically distinct cell populations are present within the same individual. Finally, SCiMS may not recapitulate the reported sex of individuals whose sex-related biological characteristics do not align with binary definitions of male or female. Therefore, while SCiMS offers a novel bioinformatic approach for predicting chromosomal sex, it is essential to acknowledge its inferences are limited to binary sex chromosome models and do not capture biological or social dimensions of sex and gender beyond chromosomal inference.

Finally, researchers should consider the ethical implications of applying SCiMS to their own or publicly available datasets, particularly when analyzing human-derived data. Sex is widely considered sensitive personal data and inadvertent disclosure of this information can have privacy implications [33, 34]. Following recommendations from a 2022 U.S. Department of Health and Human Services-sponsored report, sex should only be collected when sex is relevant to a study’s scientific aims [31]. Furthermore, host reads derived from metagenomic samples may contain other sensitive genetic information beyond sex that can reveal genetic ancestry or even individual genetic variants with potential clinical relevance [8]. While it is considered good practice to filter out host-derived reads from metagenomic datasets prior to deposition in publicly available repositories, host reads can still be present if insufficient filtering is performed [34]. Researchers are encouraged to ensure datasets they collect are deposited either in controlled access repositories (such as dbGaP) or are thoroughly filtered before deposition. Beyond technical safeguards, ethical use of inferred host metadata depends on study context and participant consent. As with any inference of sensitive host metadata, the application of SCiMS to human-derived metagenomic data should be consistent with existing ethical approvals and informed consent governing the original data collection, enabling responsible use of inferred sex information in microbiome research.

In conclusion, we developed SCiMS to fill the critical gap in accurately determining host chromosomal sex from metagenomic data, particularly at low host read coverages. Our findings demonstrate that SCiMS not only outperforms existing methods in these challenging scenarios, but also offers potential applications in sample quality control processes. While SCiMS has limitations, particularly regarding the quality of the reference genome and the sex chromosome content of the sample, it represents a valuable tool that helps scientists identify the sex of each sample. This allows them to incorporate sex as a metadata variable in their analysis or confirm metadata integrity, ultimately making microbiome research more accurate and reliable.

## Conclusions

In this study, we presented SCiMS, a scalable tool designed to predict host sex from metagenomic sequencing data. By evaluating the multinomial likelihood of observed sex-chromosome and autosomal read counts under each candidate karyotype, SCiMS infers host chromosomal sex without training data. Our benchmarks demonstrate that SCiMS performs comparably to or better than existing tools across a range of host read depths. Beyond human studies, we have shown that SCiMS effectively generalizes across diverse biological systems, including mammalian and reptile (XY) and avian (ZW) models. By providing a reliable method to recover or verify sex metadata, SCiMS inferred chromosomal sex can be incorporated into downstream statistical models, thereby enhancing the reproducibility and rigor of metagenomic studies.

## Abbreviations

FN: False Negatives
FP: False Positives
HMP: Human Microbiome Project
KDE: Kernel Density Estimation
PAR: Pseudoautosomal Region
QC: Quality Control
SCiMS: Sex Calling in Metagenomic Samples
TP: True Positives

## Declarations

### Ethics approval and consent to participate

Not applicable.

## Consent for publication

Not applicable.

## Availability of data and materials

The HMP dataset was downloaded from dbGaP Study Accession: phs000228. The Hadza dataset is available at ENA project number PREJEB49206. The Indian Human Gut Microbiome datasets is available at ENA project number PRJNA397112. The mouse cecal dataset is available at ENA project number PRJEB32890. The chicken cecal datasets are available at the ENA project number PRJEB33338 and NCBI SRA project numbers PRJNA658643 and PRJNA947933. The baboon dataset is available at ENA project number PRJNA271618. The cow dataset is available at ENA project number PRJEB31266. The pig dataset is available at ENA project number PRJNA629856. The black rhino dataset is available at ENA project number PRJNA532626. The mesquite lizard dataset is available at ENA project number PRJNA1106546.

All source code, Snakemake pipelines, Conda environment files and parameter settings to recreate the analyses in this paper are available at https://github.com/davenport-lab/SCiMS-paper. The SCiMS tool is available at https://github.com/davenport-lab/SCiMS.

## Authors’ contribution

ERD and HNT conceived of the study and wrote the manuscript. The SCiMS tool was implemented by HNT and KJK. HNT performed analyses of simulated and real metagenomic data. All authors have read and approved the final manuscript.

## Acknowledgements

We wish to thank members of the Davenport lab for their feedback and Kyle McGovern for developing an early prototype of SCiMS.

