## Supplementary Documents for "SCiMS: Sex Calling in Metagenomic Sequences"

Supplementary Materials

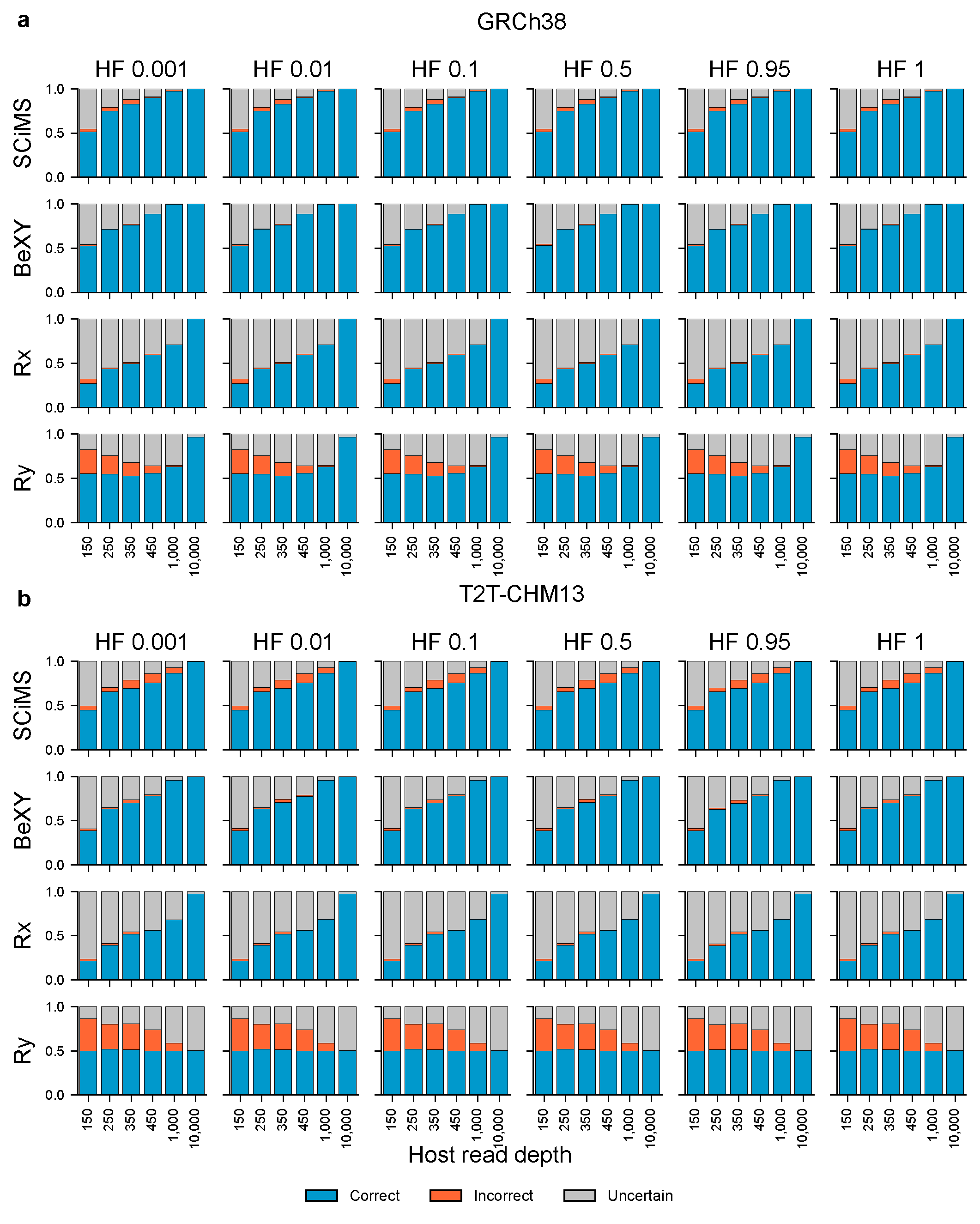

### Supplementary Figures

**Supplementary Fig. S1. Sex classification outcomes across references assemblies GRh38 and T2T-CHM13 in simulated metagenomes.** Simulated samples with known host sex generated from GRCh38 (**a**) and T2T-CHM13 (**b**) assemblies were classified by SCiMS, BeXY, Rx, and Ry. Within each panel, rows correspond to the four sex inference methods and columns represent six host read fractions (the proportion of host-derived reads in each sample, from 0.1 to 100%). Each cell shows the fraction of samples classified as correct (blue), incorrect (orange), or uncertain (grey). Host read depth was held constant within each depth bin while microbial reads were added to the target host fraction, so that host read count and host fraction vary independently.

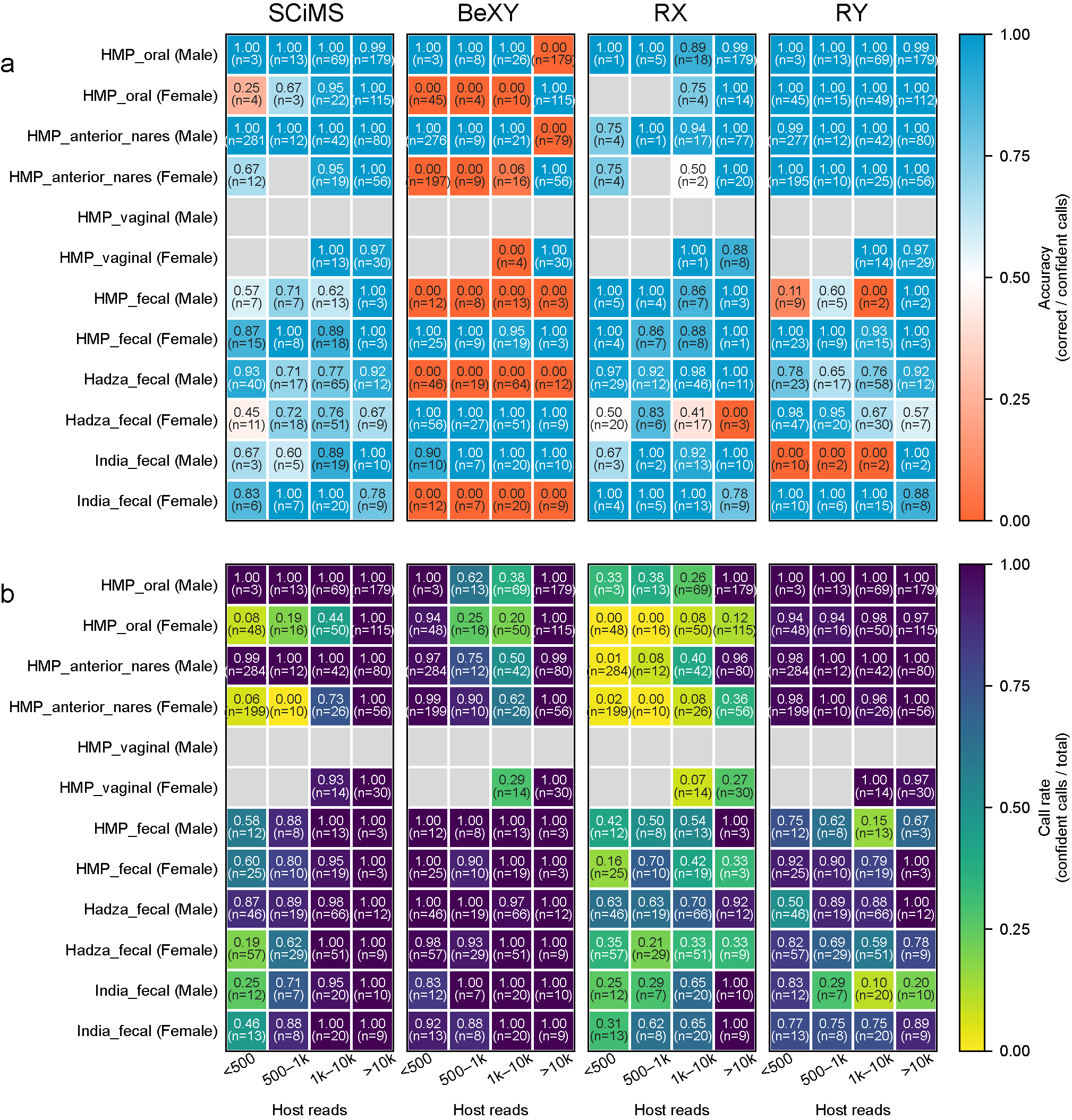

**Supplementary Fig. S2. Sex classification accuracy and call rate across human cohorts, varying by host sex and host read depth.** SCiMS, BeXY, Rx, and Ry were applied to human metagenomic samples from six cohort and body site combinations (HMP oral, anterior nares, vaginal, and stool; Hadza fecal; Indian fecal), with samples grouped by host sex and by host read depth (<500, 500-1,000, 1,000-10,000, >10,000 host reads). (**a**) Accuracy, defined as the fraction of confident calls that were correct, shown for each cohort-body site by sex (rows) at each read depth bin (columns) for the four methods. (**b**) Call rate, defined as the fraction of samples receiving a confident call (confident calls / total samples). Cell color encodes the value, and each cell is annotated with the value and the number of samples (n). Grey cells indicate group with no samples of the corresponding sex or no data available for that group.

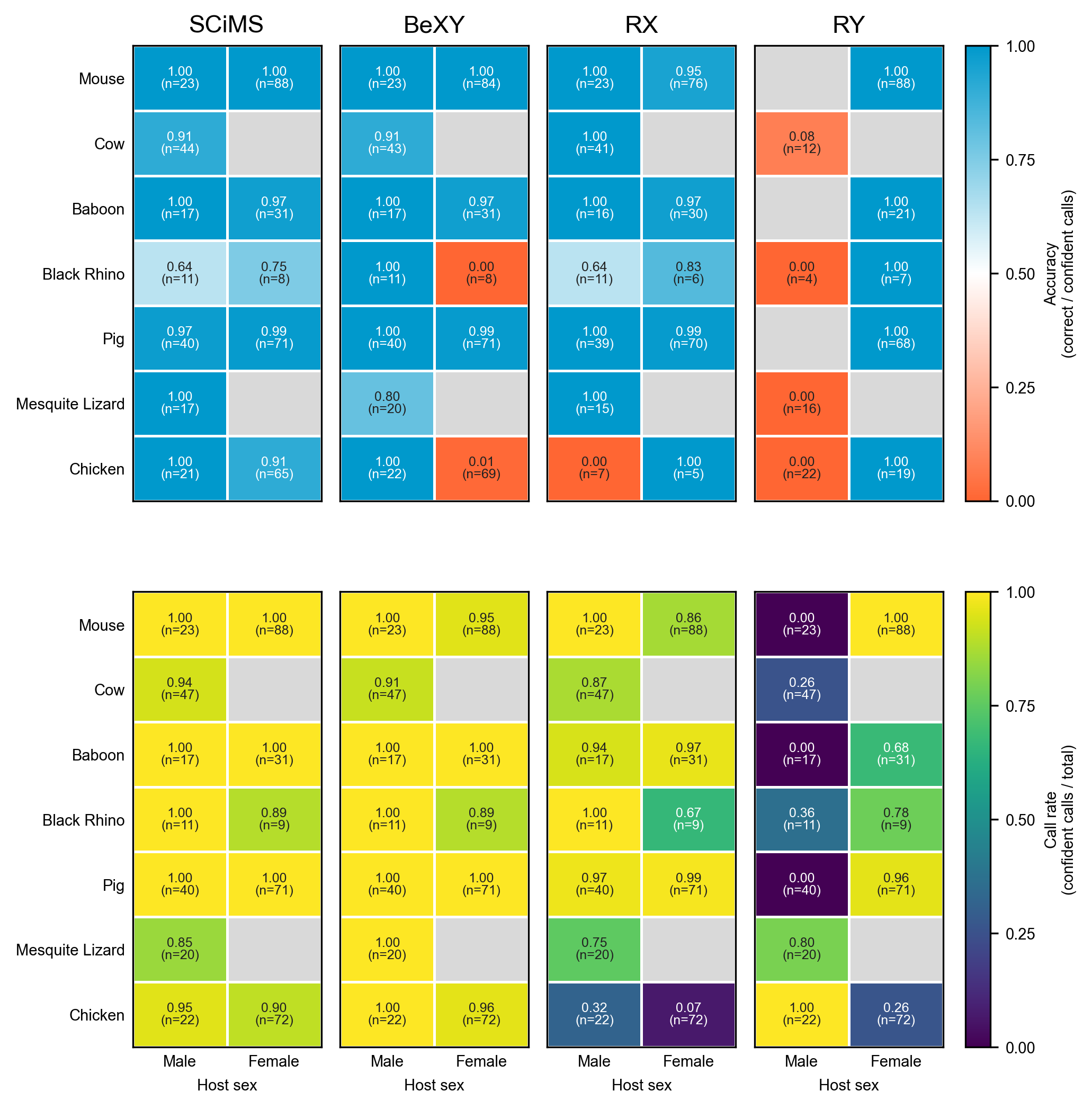
**Supplementary Fig. S3. Sex classification accuracy and call rate across animal species, varying by host sex.** SCiMS, BeXY, Rx, and Ry were applied to metagenomic samples from seven animal species (mouse, cow, baboon, black rhino, pig, mesquite lizard, and chicken), with samples grouped by host sex. (**top**) Accuracy, defined as the fraction of confident calls that were correct, shown for each animal group (rows) and host sex (columns) for the four methods. (**bottom**) Call rate, defined as the fraction of samples receiving a confident call (confident calls / total samples). Cell color encodes the value, and each cell is annotated with the value and the number of samples (n). Grey cells indicate group with no samples of the corresponding sex or no data available for that group.

Supplementary method: Mathematical Framework of SCiMS

### 1. Overview

SCiMS infers host chromosomal sex from metagenomic data using a likelihood-ratio test built on a generative model of read placement. Given a host reference genome with known scaffold lengths and sex-chromosome identifiers, the algorithm computes the posterior probability of that sample originated from a male or a female host. No training data are required because the per-chromosome read distribution expected under each sex hypothesis is derived from chromosome lengths and ploidy. This design makes SCiMS applicable to any host organisms with a heterogametic sex determination system (XY or ZW) without species-specific recalibration.

### 2. The Algorithm

2.0 Notation

Let:

$c$: the scaffolds used for sex inference (autosomes A and sex chromosomes $\{H, G\}$

$H$ : homogametic sex chromosome (X in XY systems; Z in ZW systems)

$G$ : heterogametic sex chromosome (Y in XY systems; W in ZW systems)

$L_{c}$ : length of chromosome c in base pairs

$N_{c}$ : number of reads mapped to chromosome c

$N= \sum_{c} N_{c}$ : total number of reads across the included scaffolds

$n_{c}^{S}$: ploidy of chromosome c under sex hypothesis S $\in\{ male, female\}$

$\varepsilon$ : mis-mapping rate (default: ${10}^{-4}$)

The user supplies the homogametic and heterogametic scaffolds identifiers along with the list of autosomal scaffolds to be included in the model.

#### 2.1 Ploidy under each sex hypothesis

In a heterogametic sex-determination system:

Autosome a $\in$ A: $n_{a}^{male}=n_{a}^{female}=2$

Homogametic H: $n_{H}^{male}=1, n_{H}^{female}=2$ (XY system)

$n_{H}^{male}=2, n_{H}^{female}=1$ (ZW system)

Heterogametic G: $n_{G}^{male}=1, n_{G}^{female}=0$ (XY system)

$n_{G}^{male}=0, n_{G}^{female}=1$ (ZW system)

For a chromosome with zero expected ploidy under sex S, we replace the zero with the mismapping rate $\varepsilon$ before renormalization (see Section 2.3). This prevents the log-likelihood ratio from diverging when a small number of reads spuriously map to a chromosome that should carry no reads under the true sex hypothesis.

#### 2.2 Per-chromosome read distribution

Under sex hypothesis S, each mapped read is treated as an independent draw from a categorical distribution over chromosome c. The probability of observing a read on chromosome c is proportional to the ploidy $\times$ length of c under sex S:

$$p_{c}^{S}=\frac{n_{c}^{S}\cdot L_{c}}{\sum_{j} (n_{j}^{S}\cdot L_{j})}$$

This assumes (i) uniform per-base read coverage across the included scaffolds, (ii) that mapped reads have not been preferentially filtered in a way that distinguishes autosomes from sex chromosomes, and (iii) that mappability is approximately equal across the included scaffolds. We discuss violation of these assumptions and how the mismapping rate mitigates them in section below.

Under this model, the joint distribution of read counts $\{N_{c}\}$ given total reads N is multinomial:

$$P\left( \left\{ N_{c} \right\} \right|N, S)=\frac{N!}{\prod_{c} \left( N_{c}! \right)\cdot\prod_{c} \left( p_{c}^{S} \right)\cdot\left\{ N_{c} \right\}}$$

### 2.3 Mis-mapping rate

In metagenomic data, a small fraction of reads originating from microbial genomes could spuriously map to the host reference, including to chromosomes that under a given sex hypothesis are expected to carry no reads (e.g., Y in female sample). Without correction, even a single such read would force the male-vs-female log-likelihood ratio to infinity, producing an erroneous confident call.

To address this, we replace the zero-ploidy probability with a small error rate of $\varepsilon={10}^{-4}$ before computing per-chromosome probabilities:

$n_{c}^{S}=\varepsilon$

After applying the error rate, probabilities are renormalized so that $\sum_{c} p_{c}^{s}=1$ under each sex hypothesis. The value of $\varepsilon={10}^{-4}$ corresponds to a mismapping rate of roughly one read in 10,000.

### 2.4 Likelihood ratio and posterior probability

The log-likelihood ratio comparing male and female hypotheses is:

$$\Lambda= \sum_{c} N_{c}\cdot[ log(P_{c}^{male})-log(P_{c}^{female})]$$

Under a uniform prior on sex, the posterior probability that the sample is male is given by the following logistic transformation:

$$P\left( male \right| observed data)=\frac{1}{1+exp(-\Lambda)}$$

2.5 Decision rule

SCiMS reports a confident call when the larger posterior probability exceeds a user-defined threshold $\tau$ (default $\tau=0.95$):

SCiMS predicted sex = male if $P\left( male | data \right)\geq\tau$

= female if $P\left( female | data \right)\geq\tau$

= uncertain otherwise

2.6 Output

For each input sample, SCiMS reports: the predicted sex (male, female, or uncertain); the posterior probabilities for male and female; the log-likelihood ratio; the total number of reads mapped to all scaffolds; and the number of reads mapped to each sex chromosome.
